# Bayesian estimation yields anti-Weber variability

**DOI:** 10.1101/2024.08.08.607196

**Authors:** Arthur Prat-Carrabin, Samuel J. Gershman

## Abstract

A classic result of psychophysics is that human perceptual estimates are more variable for larger magnitudes. This ‘Weber behavior’, however, has typically not been the focus of the prominent Bayesian paradigm. Here we examine the variability of a Bayesian observer, in comparison with human subjects. In two preregistered experiments, we manipulate the prior distribution and the reward function in a numerosity-estimation task. When large numerosities are more frequent or more rewarding, the Bayesian observer exhibits an ‘anti-Weber behavior’, in which larger magnitudes yield less variable responses. Human subjects exhibit a similar pattern, thus breaking a long-standing result of psychophysics. Nevertheless, subjects’ responses are best reproduced by a logarithmic encoding of magnitudes, a proposal of Fechner often regarded as accounting for Weber behavior. We thus obtain an anti-Weber behavior together with a Fechner encoding. Our results suggest that the increasing variability may be primarily due to the skewness of natural priors.

---

To usefully interact with our environment, we need internal representations of the external variables that are relevant to us. Experimental investigations in psychophysics — the study of the relation between physical, external stimuli and subjective, internal sensations — have identified since the 19th century a handful of regularities in the responses of human subjects in various perceptual tasks. In estimation tasks, a common result is that the standard deviation of magnitude estimates increases in proportion with the estimated magnitude. This finding (which has been called ‘scalar variability’) has been understood as extending to estimation tasks the prominent Weber’s law, which states that in discrimination tasks the difference in magnitude necessary to reliably distinguish two stimuli is proportional to the magnitude of the stimuli [1–4]. These observations seem to point to a general principle, that judgments about larger magnitudes come with greater variability, and hence a loss of sensitivity to magnitude differences.

The past decades have seen the development of a different line of theory regarding perceptual judgments, in which perception is conceived as resulting from a process of Bayesian inference about external stimuli, carried on the basis of imprecise (noisy) internal signals (e.g., the activity of sensory neurons), in combination with prior knowledge about the distribution of stimuli that one can expect [5–10]. The Bayesian paradigm has an appealing theoretical grounding, and it readily accounts for the pervasive variability in responses observed in estimation tasks, and for the difficulty of distinguishing two stimuli that are close in magnitude. Furthermore, with the added assumption that magnitudes are represented on a logarithmic internal scale, as proposed by Fechner [11], or that the imprecision in internal signals increases as a function of the represented stimulus, Bayesian models typically predict that greater magnitudes should result in more variable estimates [7, 9, 12]. In other words, Bayesian inference seems compatible with the psychophysical results mentioned above, and the increasing behavioral variability is seen as directly resulting from the decreasing precision of the internal signals.

In this paper, we argue that Bayesian decision theory implies important modulations of the behavioral variability even with constant precision of the internal signals, and that in some contexts the traditional psychophysical results should be contradicted. In two preregistered experiments, we show that indeed these regularities can be inverted. The noise in internal signals is an important ingredient of Bayesian models of perceptual judgments, but two other ingredients shape the responses of the Bayesian observer and their statistics: first, the prior, i.e., the relative frequencies of different stimuli; and second, the objective function, i.e., the relative importance of estimating different stimuli. The role of the prior in Bayesian inference has already been extensively studied, in particular to the extent that it impacts the *average* judgments of human subjects, in different contexts [9]. For instance, it accounts for the ‘central tendency of judgment’ [13] (a bias towards the center of the range of stimuli that are presented in a given experimental session.) But how the prior and the objective function impact the *variability* of estimates has received limited attention in the literature. Here we call ‘Weber behavior’ a behavior in which the variability of estimates increases with the magnitude (this is a less stringent requirement than that of ‘scalar variability’, which requires a linear relationship between the standard deviation of estimates and the magnitude). We show, first, that a Weber behavior may find its origin in the skewness of the prior only, or in that of the objective function; a Fechnerian encoding, or an increasing imprecision of internal signals, are not necessary assumptions. Second, we show that a simple manipulation of the prior or of the objective function, in a Bayesian observer model, yields a behavioral variability that is *opposite* to the generally accepted result that greater magnitudes entail greater variability. We call this pattern ‘anti-Weber behavior’.

We test these empirical predictions of Bayesian decision theory using numerosity-estimation tasks, where subjects are asked to judge the number of items in a briefly presented collection [14–20]. Studies of numerosity estimation have yielded results remarkably similar to those obtained in more traditional psychophysics, including Weber’s law and scalar variability [3, 4, 14, 15, 21, 22]. Together with neurobiological studies, which have revealed the tuning-curve properties of number-selective neurons [23–26], these results point to the existence of a ‘number sense’ [27], comparable to the other senses traditionally studied in psychophysics experiments, and which provides humans (and some animals) with the ability to represent approximate numerical magnitudes. Studying numerosity has the advantage of allowing us to directly ask human subjects for their estimate of a magnitude (i.e., a number) without any ambiguity about the response scale, while it is not obvious how a subject should respond when asked to estimate, for instance, the ‘loudness’ of a stimulus. We conducted two numerosity-estimation tasks: one in which we manipulated the prior, and one in which we manipulated the objective, so as to study the impact of each on the variability of estimates.

We first present these two tasks, and the specifics of the priors and of the objective functions that we utilize in different experimental conditions. We then present a model of a Bayesian observer, and we examine its behavior in the context of the two tasks; specifically, we show in which circumstances a Weber or an anti-Weber behavior is obtained. Turning to the subjects, we look at the statistics of their responses in the two tasks, and we exhibit how their behavior is qualitatively similar to that of the Bayesian observer. Finally, we fit nine variants of the Bayesian model to subjects’ data, and we compare their ability to successfully capture the behavioral patterns of the subjects. In particular, we examine in this comparison the performance of models featuring a logarithmic, Fechnerian encoding, and of models featuring a power-law encoding.

## Results

### Numerosity-estimation tasks

We now present the two numerosity-estimation tasks used in this study (more details can be found in Methods). The trial structure is the same in both tasks: the subject is presented for 500ms with a cloud of dots containing between 41 and 80 dots, and is then asked to provide, using a slider, their best estimate, *x*^, of the number of dots, *x* (Fig. 1A). In each trial the subject receives a number of points that is a decreasing linear function of their squared error, (*x*^ − *x*)^2^. At the end of the experiment, their total score is converted to a financial reward. Each experiment comprises two conditions, i.e., two blocks of 120 consecutive trials that differ in one aspect of the task: in the *‘priors experiment’*, we manipulate the relative frequencies of the numbers of dots shown, while in the *‘stakes experiment’*, we manipulate the point rewards associated with these different numbers. We now present in more details these experiments.

**Fig. 1:**
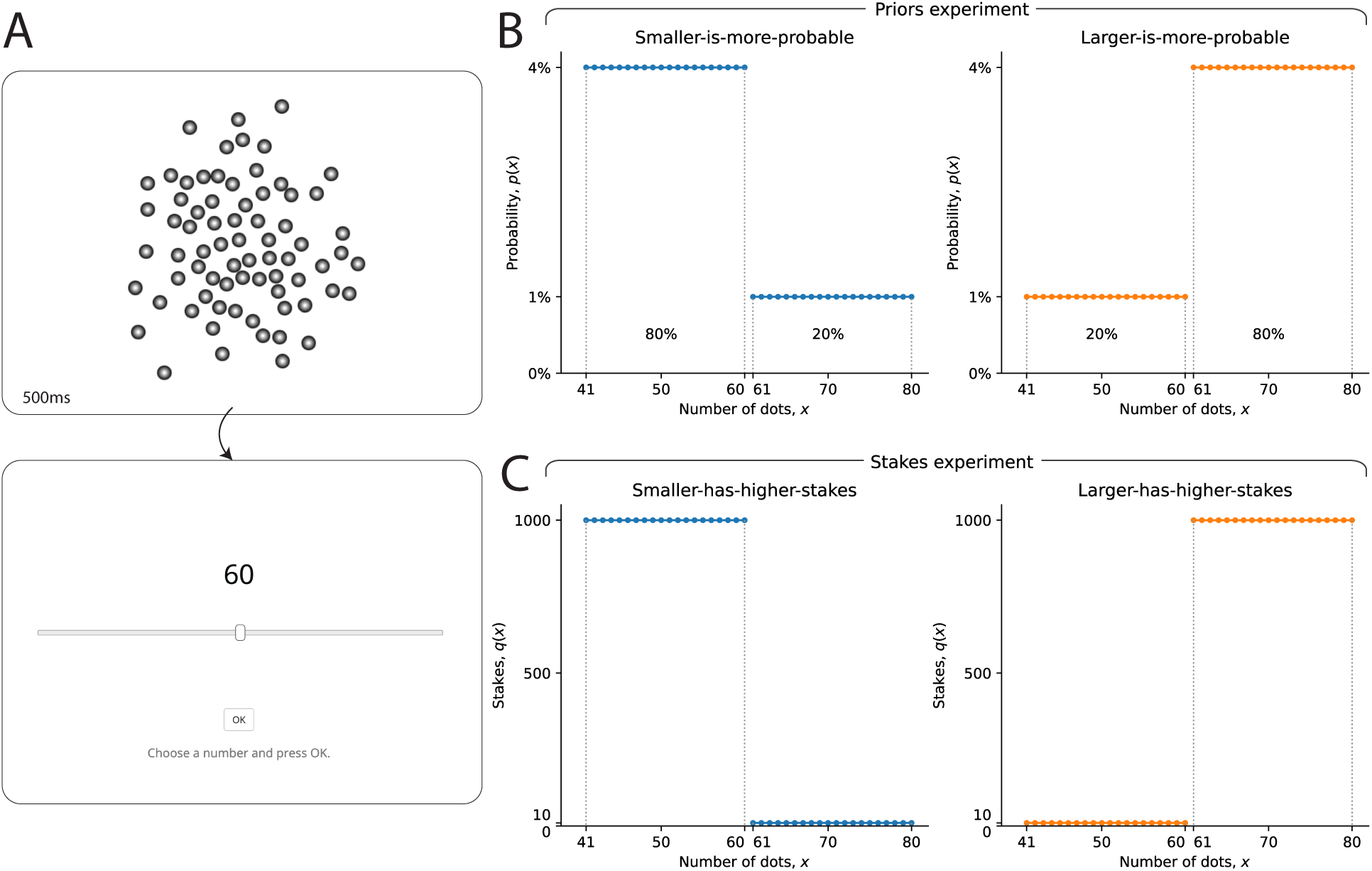
Manipulation of the priors and of the objective function in two numerosity-estimation tasks. **A.** Cloud of dots (top panel): example of visual stimulus presented to the subject for 500ms in each trial of the two numerosity-estimation tasks. Immediately after the presentation of the cloud of dots, the subject is asked to provide, using a slider (bottom panel), his or her best estimate of the number of dots in the cloud. Subjects respond at their own pace. **B.** Prior distributions from which the numbers of dots are sampled, in the two conditions of the priors experiment. In the smaller-is-more-probable condition (left panel), the probability of each number between 41 and 60 is 4%, while the probability of each number between 61 and 80 is 1%. These probabilities are inverted in the larger-is-more-probable condition (right panel). **C.** Stakes functions in the two conditions of the stake experiment. In the smaller-has-higher-stakes condition (left panel), the stakes for each number between 41 and 60 is 1000, while the stakes for each number between 61 and 80 is 10. These amounts are inverted in the larger-has-higher-stakes condition (right panel).

The two conditions of the priors experiment differ in terms of the prior, *p*(*x*), i.e., the distribution from which the number of dots is sampled on each trial. In the *‘smaller-is-more-probable’* condition, the numbers of dots between 41 and 60 are four times more probable than the numbers of dots between 61 and 80 (the total probability of the smaller numbers is 80%, while the total probability of the larger numbers is 20%; see Fig. 1B, left panel). These frequencies are inverted in the *‘larger-is-more-probable’* condition: in this condition, the larger numbers (*x* ≥ 61) are four times more probable than the smaller numbers (*x* ≤ 60; see Fig. 1B, right panel).

By contrast, in the stakes experiment, the prior is uniform in the two conditions: all the numbers (between 41 and 80) have the same probability. The two conditions of this experiment differ by the ‘stakes’, which are the maximum amount of points that a subject can get in a trial. Specifically, the stakes in each trial, *q*(*x*), are a function of the correct number of dots; and the amount of points collected in the trial is proportional to the stakes (more precisely, it is 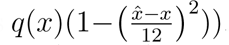. In the *‘smaller-has-higher-stakes’* condition, the stakes of the smaller numbers (*x* ≤ 60) are high, while the stakes of the larger numbers (*x* ≥ 61) are low (Fig. 1C, left panel). In the *‘larger-has-higher-stakes’* condition, conversely, the stakes of the larger numbers are high, whereas for the smaller numbers they are low (Fig. 1C, right panel). (In the priors experiment, the stakes are identical for all the numbers, in all trials.) The features of each of these conditions are explained to the subjects, in the instruction sections of the tasks. Before looking at the responses of subjects, we describe the behavior of a Bayesian observer in these two tasks.

### Patterns of variability of the Bayesian observer

We consider a Bayesian model subject for whom the presentation of a cloud containing *x* dots results in a noisy perceptual signal, *r*, on the basis of which the subject infers, using Bayes’ rule, the number of dots presented. We assume that the noisy signal is normally distributed around an increasing transformation of the correct number of dots, as

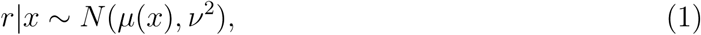

where *µ*^′^(*x*) *>* 0. The standard deviation, *ν*, parameterizes the amount of imprecision in the noisy representation of the number. In short, this model posits a mapping *µ* from the stimulus space to an internal, psychological scale, on which a stimulus *x* is represented by a noisy signal *r*, with a degree of imprecision *ν* that is independent from the stimulus.

We further assume that given an internal signal, *r*, the model subject chooses as a response the optimal estimate in the current task, *x*^*^(*r*). In our tasks, the reward is a decreasing linear function of the squared error (multiplied by the stakes, in_/_the stakes experiment), where *q*(*x*) is the stakes function (constant in the priors experiment), and *p*(*x*|*r*) is the Bayesian posterior over the numbers given the internal signal, *r* (and given the prior, *p*(*x*)). The optimal estimate (obtained by setting the derivative to zero) is a weighted average of the numbers, weighted by the prior and the likelihood (as a result of Bayes’ rule), but also by the stakes, as

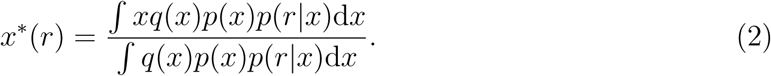

The optimal estimate *x*^*^(*r*) is a deterministic function of the noisy signal *r*; thus it is itself noisy (i.e., random), and repeated presentations of the same number *x* will result in different estimates. (Below, we also introduce motor noise in the responses of the model subject, but here, first, we assume that the selected response is the optimal estimate, and we examine the resulting behavior in this case.) The variability of the optimal estimate originates in that of the noisy internal signal, but it is also strongly shaped by the prior and by the stakes function. For instance, estimates are less variable if the product *p*(*x*)*q*(*x*) is more ‘concentrated’ (i.e., if it takes large values around some number). In a degenerate case inwhich this product is zero everywhere except at a single number, then the optimal estimate is this number, and there is no variability.

Our experimental setup provides less extreme instances of the functions *p*(*x*) and *q*(*x*), and thus we use these to illustrate how the distribution of the responses of the Bayesian observer is modulated by the prior and by the stakes function. Specifically, we consider the model subject described above, with the identity transformation *µ*(*x*) = *x*, and a noise parameter fixed to *ν* = 10 (a value close to that obtained by fitting a similar model, presented below, to subjects’ data). In the smaller-is-more-probable condition of the priors experiment, we find that when the presented numerosity is *x* = 41, the distribution of the responses of this Bayesian observer is relatively narrow, with a standard deviation just above 3. For larger numerosities, the distribution widens and its standard deviation increases (a pattern we call ‘Weber behavior’), up until *x* = 70, where the standard deviation reaches a maximum (6.5), before slightly decreasing (Fig. 2, top row, blue lines). The behavior in the smaller-has-higher-stakes condition of the stakes experiment is qualitatively similar, i.e., a Weber behavior (except that the variability does not decrease close to the upper bound, *x* = 80; Fig. 2, bottom row, blue lines). As the prior and the stakes function, *p*(*x*) and *q*(*x*), are interchangeable in the expression of the optimal estimate (Eq. 2), the differences in the distributions of the optimal estimate between the two experiments result from the specifics of these two functions, in each experiment. In particular, in the priors experiment the ratio of the probabilities of the large and small numbers is four, while in the stakes experiment the equivalent ratio, for the stakes, is 100 (Fig. 1B,C).

**Fig. 2:**
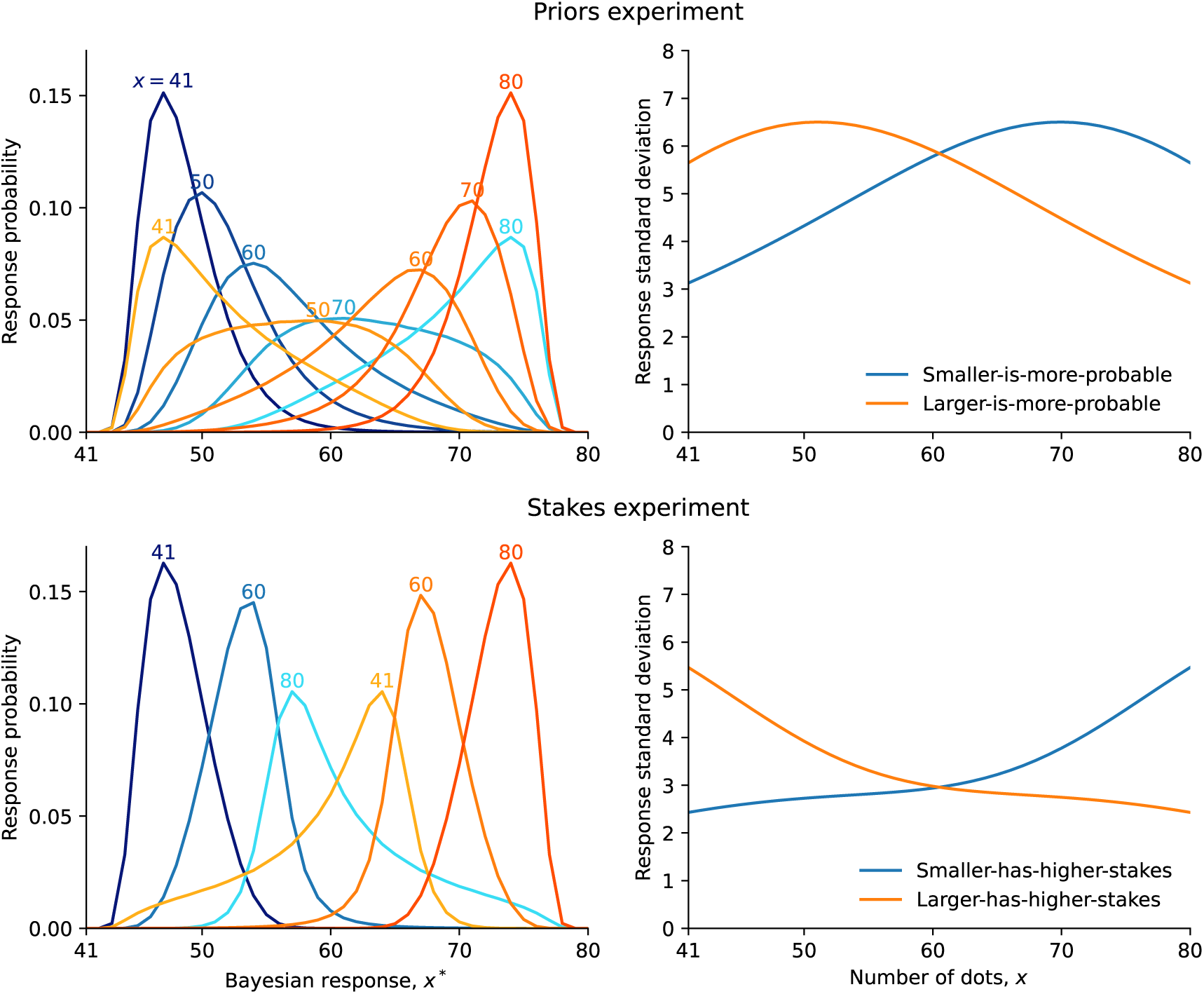
Weber and anti-Weber behavior of the Bayesian observer. Statistics of the responses of the Bayesian observer in the priors experiment (top row) and in the stakes experiment (bottom row). *Left panels:* Distribution of the responses, in the smaller-is-more-probable and the smaller-has-higher-stakes conditions (blue lines), and in the larger-is-more-probable and the larger-has-higher-stakes conditions (orange lines), for different presented numerosities (indicated above each distribution). *Right panels:* Standard deviation of the Bayesian estimate, *x*^*^, as a function of the number, *x*, in each condition.

We emphasize that here the Weber behavior we obtain does not result from a Fechnerian, logarithmic encoding of the number (as the encoding is linear, *µ*(*x*) = *x*), neither does it result from an imprecision of the internal signal that increases with the number (as the standard deviation, *ν*, is constant). This stands in contrast with the two accounts of Weber behavior most commonly found in the literature [9, 12, 14, 23, 28, 29].

Finally, the responses of the Bayesian observer in the larger-is-more-probable condition (priors experiment) and in the larger-has-higher-stakes condition (stakes experiment) mirror the behavior just described. The prior, or the stakes function, ‘attracts’ estimates towards the numbers that are more probable, or that have higher stakes; i.e., the larger numbers. This widens the distributions of estimates for small numbers, in comparison with those for large numbers, which appear narrower (Fig. 2, left panels, orange lines). Thus the variability mainly decreases as a function of the number, i.e., the Bayesian observer exhibits an ‘anti-Weber behavior’ (Fig. 2, right panels, orange lines). In sum, our model of a Bayesian observer displays a Weber behavior when small numbers have higher probabilities or higher stakes than large numbers, but it reveals an anti-Weber behavior when it is the large numbers that have higher probabilities or higher stakes. We now ask whether human subjects exhibit similar behavioral patterns.

### Weber and anti-Weber behavior of human subjects

Before examining the variability of subjects’ responses, we first look at the average responses. We find that the estimates provided by the subjects increase as a function of the presented number, in all conditions of the two experiments. However, in all conditions, subjects are biased: they tend to overestimate small numbers, and to underestimate large numbers. Such ‘central tendency of judgments’ has been reported since the beginning of the 20th century [13]. But here, we find in addition that the specifics of the subjects’ central tendency depend on the condition. In the larger-is-more-probable and larger-has-higher-stakes conditions, the average estimate for almost all numbers is significantly greater than in the smaller-is-more-probable and smaller-has-higher-stakes conditions. In other words, the response for a number depends not only on the number (as evidenced by the sensitivity of estimates to the number), but also on whether the larger numbers have high probabilities (or high stakes), in which case the responses are larger than when the larger numbers have low probabilities (or low stakes). Consequently, although in all conditions there is a number for which the bias vanishes (i.e., for which *x*^ is on average equal to *x*), this number is different depending on the condition; specifically, it is larger when large numbers are more probable, and when they have higher stakes (Fig. 3A,B).

**Fig. 3:**
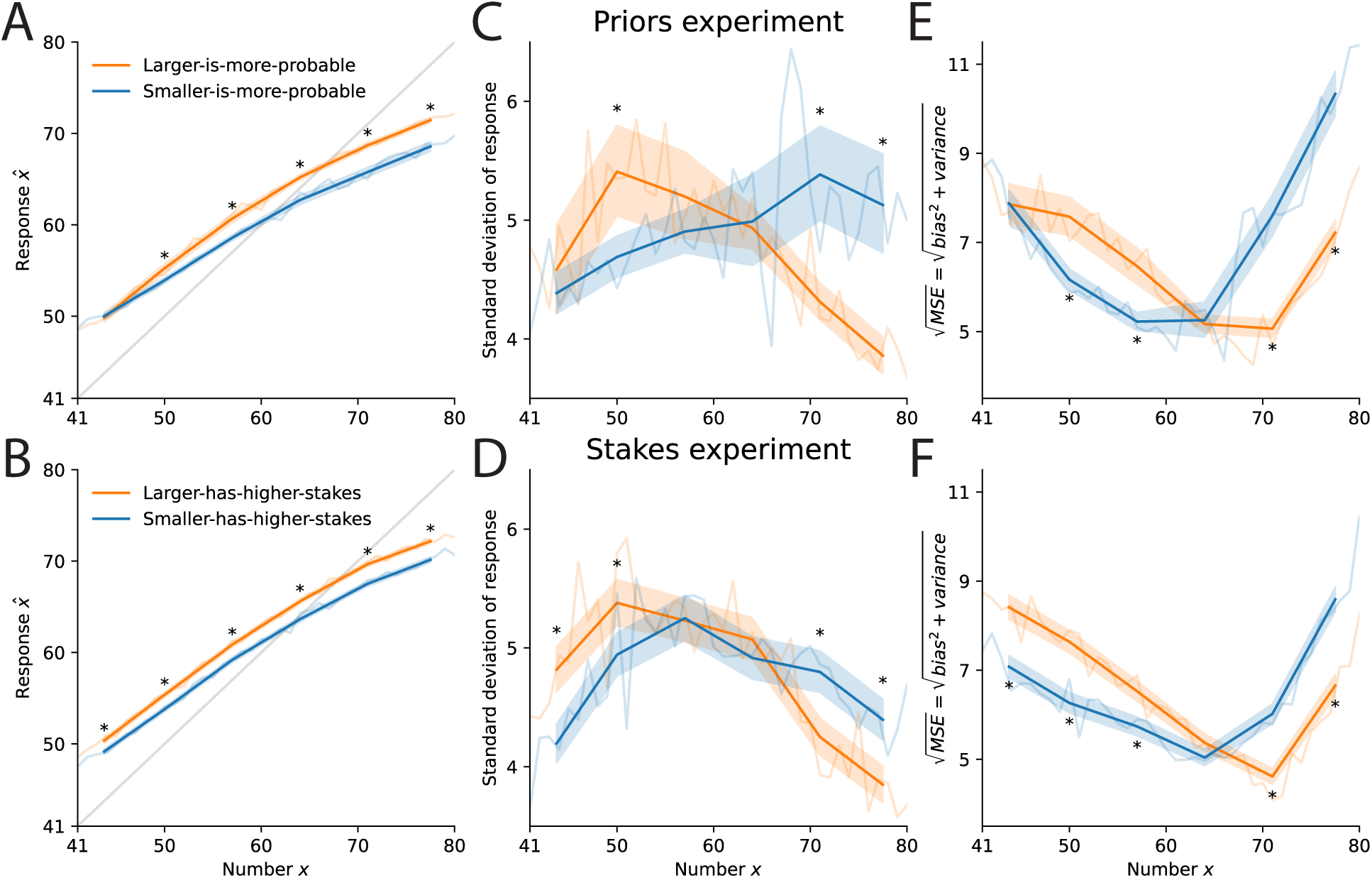
Subjects adapt the statistics of their responses to the priors and to the stakes. Subjects’ responses (A, B), standard deviations of responses (C, D), and square root of mean squared error (E, F), as a function of the presented number, in the priors experiment (A, C, E) and in the stakes experiment (B, D, F), with data grouped in six bins of the number (dark lines), or not (light-colored lines). Shaded areas show the 5%-95% credible intervals, and stars indicate Bayesian p-values < 0.005 across conditions (see Methods).

We turn to the standard deviation of subjects’ responses, in each condition of the two experiments. In the smaller-is-more-probable condition of the priors experiment, the standard deviation is an increasing function of the presented number, for numbers up to around 70, and it slightly decreases for larger numbers (Fig. 3C, blue line). We conclude that over most of the range of numbers presented, the subjects’ variability is consistent with a Weber behavior, as was found in other studies [4, 15, 28, 30, 31]. Turning to the larger-is-more-probable condition, we find that the variability decreases with the presented numbers (except near the lower boundary, where it increases), and thus that it presents a pattern opposite to that obtained in the smaller-is-more-probable condition (Fig. 3C, orange line). In other words, the behavior exhibited by subjects in this condition is of the anti-Weber kind. As a result, the variability of responses to the large numbers is significantly lower, when these are more frequent, than when the smaller numbers instead are more frequent. In the stakes experiment, a similar pattern emerges, and in particular we find again an anti-Weber behavior of subjects, in the larger-has-higher-stakes condition (Fig. 3D, orange line). Overall, the behavioral patterns of the subjects are qualitatively consistent with those of the Bayesian observer (compare Fig. 3C,D to Fig. 2, right panels).

They are also consistent with the predictions that we had included in our preregistrations of the experiments (see Methods). For the priors experiment, our prediction was that the variance of estimates would be lower when the probability of the presented number is higher. We thus conducted two Levene’s tests of equality of the variances between the smaller-is-more-probable and the larger-is-more-probable conditions, as detailed in the preregistration: one for the small numbers (*x* ≤ 60) and one for the large numbers (*x* ≥ 61). The p-values for the two tests were 0.002 and 4e-24 (*F* (1, 8914) = 9.51 and *F* (1, 8842) = 103.26), and in both cases our prediction regarding the sign of the difference was correct. For the stakes experiment, our prediction was that the variance of estimates would be lower when the stakes are higher. We thus similarly conducted two Levene’s tests, whose p-values were 1.2e-5 and 7.5e-10 (*F* (1, 13904) = 19.20 and *F* (1, 13932) = 37.93), and the signs of the differences were also as predicted. In short, all our preregistered predictions were verified.

To summarize, we find bias and variability in subjects’ responses, and we find that they both depend on the shapes of the prior and of the objective function. In the Bayesian model, the subject takes into account these two functions in order to provide a response that minimizes a loss function. Hence we investigate whether subjects seem to modulate the statistics of their responses so as to minimize their loss. Looking at the square root of their mean squared error (MSE), which essentially combines the errors brought about by the bias and by the variability, we find that it is not a constant function of the presented number. The MSE, instead, adopts a U shape, and crucially its minimum is reached at a different number in the different conditions: specifically, in the larger-is-more-probable and the larger-has-higher-stakes conditions, it is minimized at a larger number than in the other two conditions (Fig. 3E,F). In other words, the subjects minimize their errors for the presented numbers that “matter” more, either because they are more frequent, or because they have higher stakes in the estimation task. While a traditional Weber behavior implies that larger magnitudes should lead to larger errors in estimation, our results indicate that human subjects can make, conversely, smaller errors for large magnitudes, if it is warranted by the prior or by the objective function.

### Best-fitting Bayesian model

Our main goal in this study is to investigate the implications of Bayesian inference with respect to behavioral variability, and examine in comparison the variability of human subjects. In the previous sections, we have shown that subjects, like our model of a Bayesian observer, indeed exhibit Weber and anti-Weber behavior, depending on the experimental condition. Here, we seek to describe in some more detail the ability of the Bayesian approach to capture the patterns that we have identified in the behavioral data. We thus fit several Bayesian models to the responses of the subjects (by maximizing their likelihoods). These models are variants of the model we have presented. First, they all include some noise in the selection of the response (‘motor noise’): given the optimal estimate *x*^*^ (derived through Eq. 2), the response *x*^ is sampled from a Gaussian distribution centered on *x*^*^, with standard deviation *σ*, and ‘clipped’ (or ‘rectified’) to remain in the legal response interval (41 to 80), i.e., responses that would be outside of this range are replaced instead by the corresponding extreme value, 41 or 80 (in our investigations, we have found that this better accounts for the data than a ‘truncated’, renormalized Gaussian distribution).

Second, although we have seen that a linear encoding (*µ*(*x*) = *x*) yields a Weber behavior in the smaller-is-more-probable and smaller-has-higher-stakes conditions, the prominence of the Fechnerian view prompts us to examine a logarithmic encoding (*µ*(*x*) = log *x*). Thus we implement both encodings, in variants of the model that we label accordingly as ‘linear’ and ‘Fechner’ models. Another non-linear encoding, prominent in magnitude estimation studies, is the power law [32], in which the magnitude is raised to an exponent *a*, as *µ*(*x*) = *x*^a^. Equivalently we can choose any affine transformation of this encoding (as the resulting distribution of Bayesian estimates would be the same). We thus implement ‘power-law’ variants of the model, with the encoding function *µ*(*x*) = (*x*^a^ − 1)*/a*, for *a* ≠ 0s, which we extend, for *a* = 0, to its limit lim_a→0_(*x*^a^ − 1)*/a* = log *x*. With this specification, the power-law variants of the models nest the Fechner variants (with *a* = 0) and the linear variants (with *a* = 1), and values of *a* lower than 1 indicate a compression of the magnitude space in the encoding.

Finally, we surmise that the subjects may not perfectly learn the prior and the stakes function in each condition, although these are fully described in the instructions. The conditions of the experiments are characterized by the ratio between the prior probabilities, or between the stakes, of the large vs. the small numbers (Fig. 1B,C). Thus we implement variants of the model, which we label ‘subjective’, in which we allow the model subject to derive its estimate on the basis of a subjective ratio that may deviate from the correct value. In ‘subjective-symmetric’ models, the subjective ratio is the same in the two conditions of each experiment (as is actually the case, in the experiments), while in ‘subjective-asymmetric’ models the subjective ratio is allowed to be different: for instance, in the stakes experiment, the model subject may believe that large numbers in the larger-has-higher-stakes condition have proportionally greater stakes than the small numbers in the smaller-has-higher-stakes condition. By contrast, the correct ratio is used in variants of the model that we call ‘correct’. We thus obtain nine models, specified by the choice of the encoding, ‘linear’, ‘Fechner’, or ‘power-law’, and by whether the priors and the stakes used in derivations are ‘correct’, ‘subjective-symmetric’ or ‘subjective-asymmetric’. The linear and Fechner correct models have two parameters, *ν* and *σ*, that determine the imprecision in the internal signal and in the choice of response, respectively; and the power-law correct model has a third parameter, the exponent *a*. The ‘subjective-symmetric’ counterparts of these models have an additional parameter, the subjective ratio applied in both conditions, which we denote by *ρ*; and the ‘subjective-asymmetric’ models have two additional parameters, *ρ*_s_ and *ρ*_l_, the two ratios used in the two conditions. (Specifically, *ρ*_s_ for the smaller-has-higher-stakes and the smaller-is-more-probable conditions, and *ρ*_l_ for the other two; the correct values would be *ρ*_s_ = *ρ*_l_ = 4 in the priors experiment, and *ρ*_s_ = *ρ*_l_ = 100 in the stakes experiment.) For each model, we compute its Bayesian Information Criterion (BIC), a measure of fit that penalizes additional parameters [33]. We find that the ‘subjective’ models improve the BICs by a sizable amount, thus warranting the additional parameters (Table 1).

**Table 1:** The Fechner, subjective model is prevalent among subjects in both experiments. Each row in the first part of the table corresponds to a model; the second part corresponds to models grouped by their type of encoding; the third part groups models according to the beliefs about the ratios characterizing each condition. BIC: Bayesian Information Criteria (lower is better). *p* and PXP: expected probability of each model or group of models, and their protected exceedance probability (see Methods). For PXPs, an empty cell means PXP < 0.0015 and 1^*^ means PXP > 0.9985.

| Model |  | Priors experiment |  |  | Stakes experiment |  |  |
| --- | --- | --- | --- | --- | --- | --- | --- |
| | | BIC | $p$ | PXP | BIC | $p$ | PXP |
| Fechner | Subjective (Asym.) | 107,192 | 0.31 | 0.54 | 171,065 | 0.37 | 0.47 |
| Power-law | Subjective (Asym.) | 107,308 | 0.06 |  | 171,164 | 0.10 |  |
| Linear | Subjective (Asym.) | 107,266 | 0.24 | 0.13 | 171,091 | 0.37 | 0.53 |
| Fechner | Subjective (Sym.) | 108,793 | 0.28 | 0.33 | 176,312 | 0.09 |  |
| Power-law | Subjective (Sym.) | 108,945 | 0.02 |  | 176,085 | 0.03 |  |
| Linear | Subjective (Sym.) | 109,015 | 0.03 |  | 176,561 | 0.01 |  |
| Fechner | Correct | 112,298 | 0.01 |  | 183,655 | 0.01 |  |
| Power-law | Correct | 112,460 | 0.02 |  | 183,796 | 0.01 |  |
| Linear | Correct | 112,475 | 0.02 |  | 183,649 | 0.01 |  |
| Fechner |  |  | 0.60 | 1* |  | 0.47 | 0.82 |
| Power-law |  |  | 0.10 |  |  | 0.14 |  |
| Linear |  |  | 0.30 |  |  | 0.39 | 0.18 |
|  | Subjective (Asym.) |  | 0.61 | 0.99 |  | 0.84 | 1* |
|  | Subjective (Sym.) |  | 0.34 | 0.01 |  | 0.13 |  |
|  | Correct |  | 0.06 |  |  | 0.03 |  |

We use random-effects Bayesian model selection to compare models [34, 35]. In this procedure, the behavior of each subject is treated as a random draw from a distribution over the models, which is estimated using the data. For each model we report the expected probability, *p*, which is an estimate of the fraction of the population that is best captured by the model. Together, the Fechner models represent in both experiment a relative majority, with a total expected probability of 60% and 47% in the priors experiment and in the stakes experiment, respectively, compared to 30% and 39%, respectively, for the linear models, and 10% and 14%, respectively, for the power-law models (Table 1). We also report the ‘protected exceedance probability’ (PXP), a conservative estimate of the probability that the model is the most prevalent in the population. The PXPs substantiate the relative prevalence of the Fechner models, with values close to 100% in the priors experiment and 82% in the stakes experiment. Moreover, the median exponent parameter *a* of the power-law model (in its subjective-asymmetric variant) is 0.18 in the priors experiment and 0.24 in the stakes experiment, and the best-fitting exponent is lower than 1 for 76% and 72% of subjects, respectively. Overall, we conclude that a compressive encoding of the numerosities is dominant in the population, with a significant fraction of subjects well captured by Fechner models, thus supporting the hypothesis that a logarithmic encoding underlies the representation of numerical magnitudes [23, 29].

Although across the two conditions of each experiment the two priors (or the two stakes functions) are symmetric to each other, model fitting suggests that the subjects do not learn them symmetrically, as evidenced by the lower BICs and the greater expected probabilities and PXPs of the ‘subjective-asymmetric’ models (Table 1). Subjects in the two experiments seem more sensitive to the conditions in which larger numbers are more important (i.e., *ρ*_l_ *> ρ*_s_ for most subjects, in both experiments). With the ‘Fechner, subjective-asymmetric’ model (whose BIC is the lowest in both experiments), the ratio of the two parameters, *ρ*_l_*/ρ*_s_, has a median (across subjects) of 1.34 in the priors experiment, and of 2.47 in the stakes experiment (instead of 1 in the correct, symmetric case), with a large inter-individual variability (inter-quantile range (IQR): 0.53-5.50 and 0.61-21.1 in the priors and stakes experiment, respectively). This suggests that for most subjects large numbers in the larger-has-higher-stakes and larger-is-more-probable conditions have greater stakes and higher probabilities, respectively, than the small numbers in the smaller-has-higher-stakes and smaller-is-more-probable conditions.

Furthermore, the best-fitting ratios *ρ*_s_ and *ρ*_l_ suggest a strong attenuation in how most subjects incorporated the difference between the probabilities or the stakes of the two halves of the magnitude space. Indeed the median ratio *ρ*_s_ is close to 1 in the stakes experiment (median: 1.01, IQR: 0.24-2.44), and also to a lesser extent in the priors experiment (median: 1.16, IQR: 0.49-10.8), implying that in the smaller-has-higher-stakes condition of the stakes experiment, almost half of the subjects consider that small numbers, in fact, have *lower* stakes than the large numbers — although not as low as in the larger-has-higher-stakes condition. This would account for the fact that the subjects’ variability in the smaller-has-higher-stakes condition reaches its maximum where numbers have high stakes, in the lower half of the interval (and their MSE reaches a minimum in the larger half; Fig. 3D,F). We surmise that this may stem from the ecological relevance of large quantities, typically associated with higher stakes in real-world contexts. This question however is beyond the scope of our study; here we emphasize that our manipulation of the stakes function yields a behavioral effect in the expected direction. As for the ratio *ρ*_l_, as mentioned, it is generally larger than *ρ*_s_ (stakes experiment, median: 2.93, IQR: 1.09-9.45; priors experiment, median: 2.14, IQR: 0.97-7.55). Aside from these ratios, the noise parameters are stable across the two experiments: the median best-fitting value of *ν* is 0.14 in both experiments (IQR, priors experiment: 0.11-0.18; stakes experiment: 0.11-0.17), and the median best-fitting value of *σ* is 2.05 (1.17-3.04) in the priors experiment and 2.34 (1.54-3.04) in the stakes experiment.

We simulate the Fechner model and the linear one (both in their ‘subjective-asymmetric’ variants) and examine the statistics of their responses. They provide a good qualitative match with subjects’ responses. The models reproduce the central tendency of estimates, and the way it is modulated by the condition (with larger responses in the larger-is-more-probable and larger-has-higher-stakes conditions; Fig. 4A,B). The standard deviations of the models’ responses differ in the two conditions of each experiment, in a way similar to that of the subjects. In particular, in the larger-is-more-probable and larger-has-higher-stakes conditions (orange lines) the standard deviation for large numbers is lower than that in the other two conditions (blue lines), and for small numbers it is higher than that in the other conditions; the standard deviation reaches its maximum at a number that is smaller than the number at which the maximum is reached in the other conditions, with a more modest difference in the stakes experiment (Fig. 4C,D). The same patterns are found in the behavioral data (Fig. 3C,D). We note in addition that the standard deviations of the Fechner model, in the larger-is-more-probable and larger-has-higher-stakes conditions, decrease with the number over most of the range of numbers, although the logarithmic encoding in this model has precisely been proposed as an account of Weber behaviors. Finally, the MSEs have a U shape similar to the subjects’, also with a minimum reached at a larger number in the larger-is-more-probable and larger-has-higher-stakes conditions (Fig. 4E,F).

**Fig. 4:**
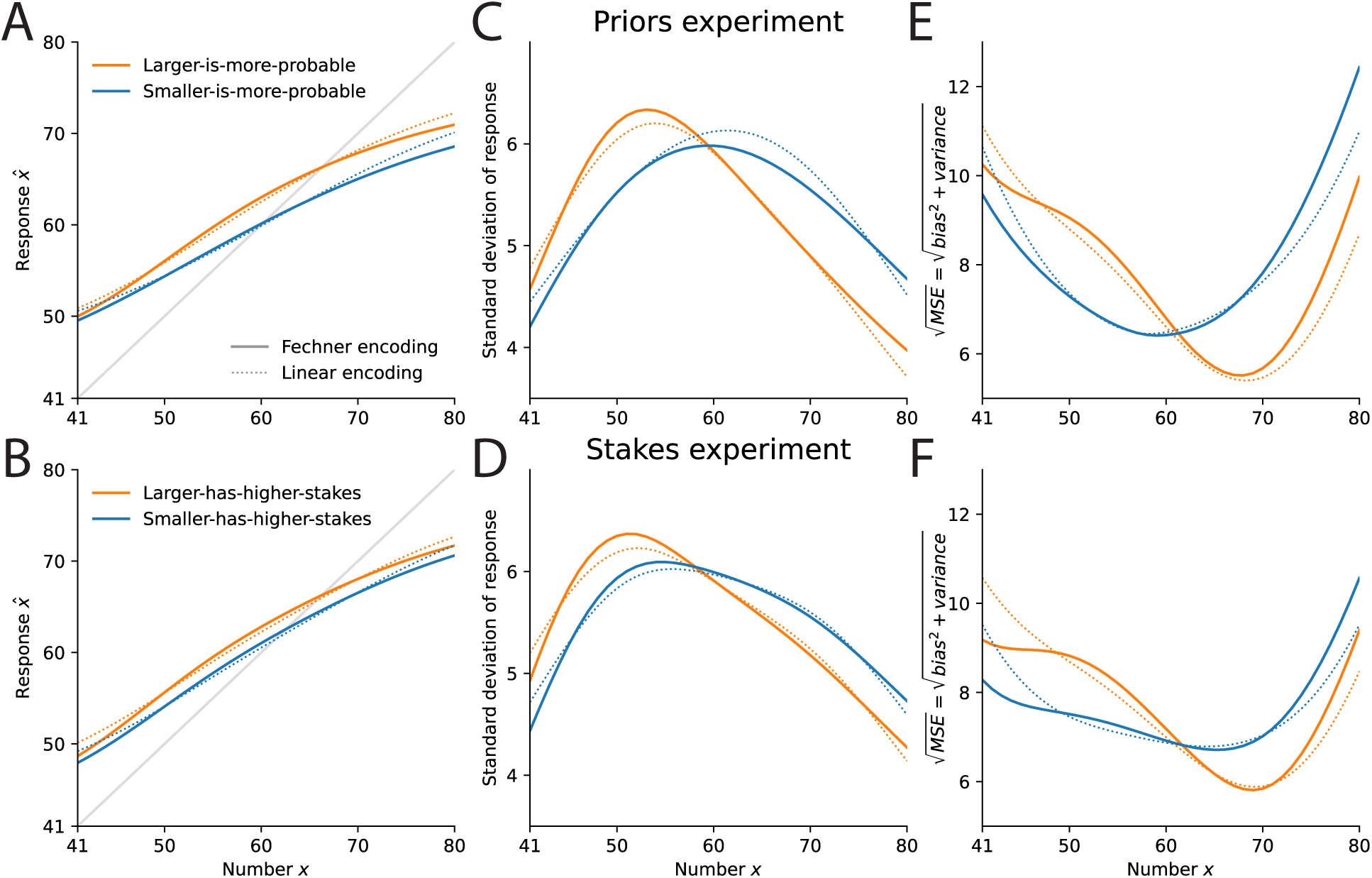
The Bayesian models reproduce subjects’ behavior. Model subjects’ responses (A, B), standard deviations of responses (C, D), and square root of mean squared error (E, F), as a function of the presented number, in the priors experiment (A, C, E) and in the stakes experiment (B, D, F), with the Fechner, subjective model (solid lines) and with the linear, subjective model (dotted lines). Compare to subjects’ behavior in Fig. 3.

## Discussion

We compared the behaviors of human subjects and Bayesian models in two numerosity-estimation tasks. Across the conditions of these tasks, small and large numbers differ either by the relative frequency in which they appear (e.g., in one condition, small numbers are more frequent), or by the reward associated with their estimation (e.g., in one condition, estimating correctly a small number brings more points than estimating correctly a large number; Fig. 1). The Bayesian observer takes into account these differences in the relative ‘importance’ of small and large numbers, and this influences the statistics of its responses. In particular, we show how it yields a Weber behavior, when small numbers are more frequent or more rewarding, and conversely an anti-Weber behavior, when large numbers instead are more frequent or more rewarding (Fig. 2). The examination of subjects’ responses reveal similar patterns (Fig. 3). Notably, we find a decreased variability for larger magnitudes, a finding that directly conflicts with traditional results of psychophysics. In short, our results suggest that the behavior of subjects is consistent with a model of Bayesian inference, and that their variability exhibit a Weber behavior only when a Weber behavior is indeed predicted by the Bayesian model.

A crucial feature of the distributions that we use in the priors experiment is that they are skewed. In particular, in the smaller-is-more-probable condition, the prior is right-tailed, i.e., the mass of the distribution is concentrated on the smaller numbers. A similar skewness characterizes the empirical distributions of numbers observed in various contexts, which have been approximated by power-law distributions [36–40]; in turn, power laws have been used to model priors over numerosities [4, 41] (other studies have posited log-normal priors, which are similarly right-tailed [31, 42–44]). Our results suggest that the shape of the prior impacts the variability of estimates, and in particular that this skewness may participate in the emergence of a Weber behavior. Indeed with the right-tailed prior used in our experiment the Bayesian observer exhibits a Weber behavior (Fig. 2), and thus this behavior may more generally originate in the skewness of natural magnitude distributions. To complement the analysis presented in the Results section, we look at the variability of the Bayesian observer equipped with a linear encoding (*µ*(*x*) = *x*), and a power-law prior with exponent 2 (*p*(*x*) ∝ 1*/x*^2^; this is the exponent found in most studies on the natural frequencies of numbers [36–38]). We simulate this model subject with three different degrees of internal noise: *ν* = 5, 10, and 20. The resulting standard deviation of estimates is an increasing function of the number *x*, up to a maximum that is reached for *x* ≈ 4*ν* (above that, the standard deviations plateaus at a value close to *ν*; Fig. 5). In other words, in any experiment in which such a Bayesian observer is asked to estimate numbers that are below four times the magnitude of its imprecision, the resulting behavior will exhibit approximate scalar variability — although the encoding itself is not more precise about some numbers than others (the encoding Fisher information, a measure of the encoding precision, is in this model constant and equal to 1*/ν*^2^).

**Fig. 5:**
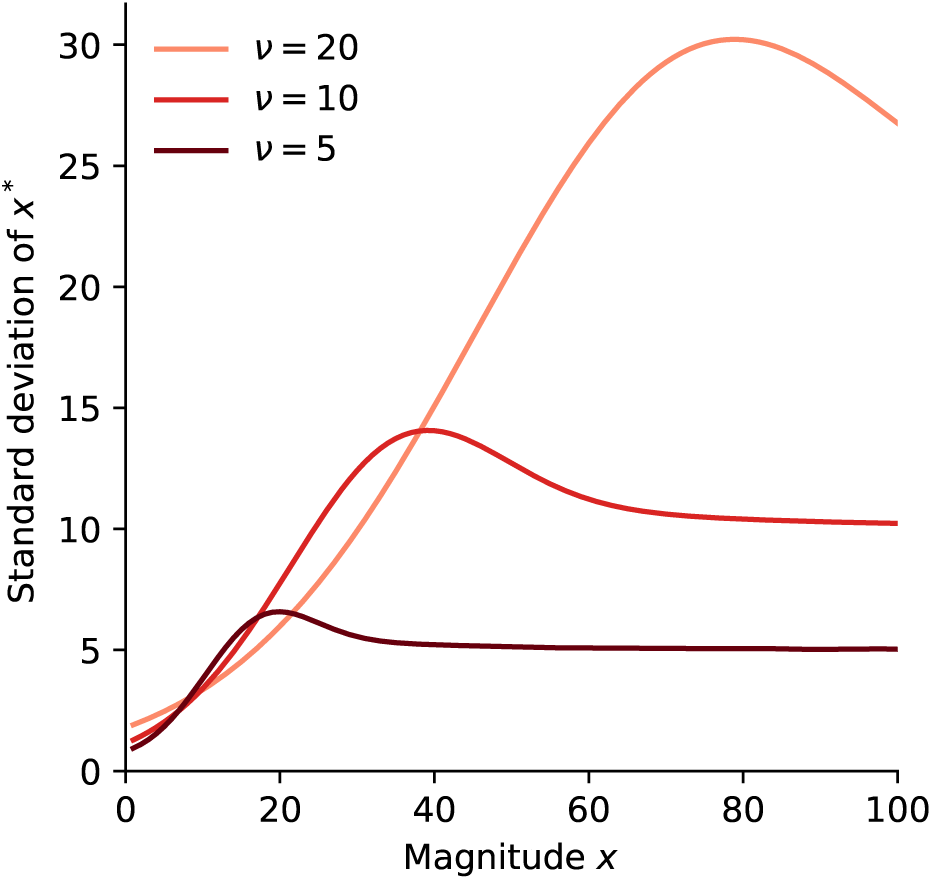
Weber behavior obtained with a linear encoding and a power-law prior. Standard deviation, as a function of the magnitude *x*, of the estimates *x*^*^ of the Bayesian observer with linear encoding (*µ*(*x*) = *x*), when the prior follows a power-law (*p*(*x*) ∝ 1*/x*^2^), and with three different values of the internal noise parameter: *ν* = 5, 10, and 20.

This account of Weber behaviors with a linear encoding does not however preclude the possibility of a logarithmic, Fechnerian encoding (*µ*(*x*) = log *x*), and in fact we find that the most prevalent model features such a logarithmic encoding. (Here we note that identifying the encoding was however not the first aim of our study. Had it been our goal, an estimation task with a uniform prior would presumably have been a more appropriate choice.) The logarithmic encoding is more precise about small numbers than about large numbers (its Fisher information decreases with the number *x*, as (*µ*^′^(*x*)*/ν*)^2^ = (*νx*)^−2^); this ‘diminishing marginal precision’ leads to a greater variability of estimates for larger numbers. Supporting the hypothesis of a logarithmic internal scale, neurophysiological investigations have exhibited numerosity-selective neurons, whose tuning curves are best described on a logarithmic scale [14, 24, 25]. Why should the brain represent numerosities on such a nonlinearly compressed scale? A possibility is that this results from an optimal adaptation to the distribution of numbers that one will need to represent, under limited resources available for representation. This idea is formalized in models of efficient coding, which typically predict that the encoding Fisher information should be proportional to the prior raised to some exponent. Specifically which exponent best reflects the constraints of the neural encoding remains unclear, and seems to depend on the encoding objective (typical values are between 1/2 and 2 [20, 45–49]). But if the distribution of numerosities that one typically encounters is well approximated by a power law, as suggested by the studies mentioned above, then the Fisher information predicted by efficient-coding models is a (negative) power of the represented magnitude. The precise exponent will depend on the specifics of the prior and of the efficient-coding model, but with a power-law prior with exponent 2 as assumed above and an efficient-coding exponent of 1, the optimal Fisher information is proportional to 1*/x*^2^; with our encoding model (Eq. 1) this would be achieved with a logarithmic encoding (*µ*(*x*) = log *x*). With different exponents, one obtains a power-law encoding of the kind we have implemented here, and which seems to also yield a reasonably good account of the data (Table 1). Ref. [37] presents a similar, ‘rational-analysis’ derivation of such non-linear encodings (see also Ref. [50]); and Ref. [51] presents an efficient-coding model under two priors similar to ours, which they examine in the context of risky choices. (In a different domain, visual working memory, tests of resource-rationality under manipulations of the reward have produced conflicting results [52, 53]). More generally, if larger magnitudes are less frequent, then under efficient coding they should be represented with decreasing precision. Alternatively, the apparent logarithmic encoding of numerosity could emerge from the properties of the brain’s processing of visual input, at least when numerosities are presented as visual arrays of multiple items; for instance neural-network models of the visual stream, which are not trained for numerosity discrimination, nonetheless exhibit numerosity-tuned responses [54–56].

Regardless of the origins of the logarithmic encoding, it is often regarded, since Fechner’s proposal, as accounting for Weber behaviors. Our results suggest that the logarithmic encoding may in fact be neither necessary nor sufficient to account for Weber behaviors. Indeed, we have shown, first, that a model with linear encoding can yield a Weber behavior (Figs. 2, 5). Second, we have seen that subjects’ responses are overall best captured by a Fechner-encoding model. One might thus think that this should imply that they exhibit a Weber behavior, but instead, they adopt an anti-Weber behavior in the larger-is-more-probable and the larger-has-higher-stakes conditions. Moreover, in these conditions, the variability of this model also decreases with the magnitude over a large part of the magnitudes’ interval, although this model features a Fechner encoding (Fig. 4C,D). The prevalence of Weber behaviors, thus, may result from the skewness of the (possibly subjective) priors used by subjects in psychophysics experiments. Many magnitudes seem indeed to follow power laws and other skewed distributions [57–62].

Other kinds of encoding have been proposed to account for Weber behaviors. In another prominent model, the encoding is linear, but the scale of noise is a linear function of the encoded magnitude [28, 63–65]. Reference [65] generalizes this approach by showing that many encoding schemes yield Weber’s law, provided that their Fisher information is inversely related to the square of the magnitude. Here our primary goal was to examine the behavior of subjects under different priors and different reward functions; we leave the investigation of which of these alternative encodings best capture the behavioral data to future studies.

We note that the term ‘Weber’s law’ primarily refers to an empirical property of just-noticeable differences in discrimination tasks. This should be distinguished from several other notions. First, it relates to threshold discrimination (i.e., difference detection), and in a strict sense it is unrelated to suprathreshold discrimination. Second, it is different from ‘scalar variability’, which pertains to estimation tasks. Third, Weber’s law is different from Fechner’s law: most notably, the latter introduces a notion of subjective sensation magnitude that is entirely absent in the former. In practice, however, ‘Weber’s law’ often refers to the idea that the subjective dissimilarity between two stimuli is determined by the ratio of their magnitudes, a proposal that has been dubbed the ‘W-principle’, or the ‘Weber principle’— in an effort, precisely, to distinguish it from Weber’s law [66, 67]. As for scalar variability, from the start it was associated with Weber’s law [1, 2], and it is telling that several studies which involve estimation tasks examine the ‘Weber fraction’ of subjects, defined in this context as the ratio of the standard deviation to the mean of estimates [3, 7]. A common thread in these various concepts is the diminishing sensitivity of perceptual judgments with the perceived magnitudes, and it is a similar notion that we have kept in our definition of a ‘Weber behavior’, whereby the variability of estimates increases with the magnitudes presented. This has made straightforward the introduction of ‘anti-Weber’ behaviors, readily defined as a decrease of the variability with the magnitude. Both are properties of responses obtained in estimation tasks, that do not rely on a putative subjective sensation magnitude. An important question is whether similar manipulations of the prior and of the stakes in a discrimination task would exhibit an increasing ability to discriminate stimuli of increasing magnitudes, when large magnitudes are more frequent or more rewarding. We leave the examination of this question to future studies.

In all conditions of our experiments we find that large numbers are on average underestimated, and small numbers are overestimated (Fig. 3A,B). This pattern is also found in the responses of the Bayesian model (Fig. 4A,B), and indeed the ability of Bayesian models to capture such ‘central tendency of judgments’ [13] has already been noted [9]. The central tendency has been obtained with various kinds of stimuli [7, 9, 68], including numerosities [16, 69]. In many of these studies, three different ranges of magnitudes are typically used in different experimental conditions, and the magnitudes in a given condition are sampled from a uniform distribution over the corresponding range. The subjects’ estimates, in each condition, are then shown to be biased towards the center of the range; consequently, the same magnitude results in different estimates, depending on the current range. Thus these studies manipulate the range of the prior, but not its shape. Here, we do not manipulate the range of the prior, which is [41, 80] in all the conditions, and instead we manipulate its shape (in the priors experiment). We find that the central tendency is modulated by the shape of the prior, in a way that is readily accounted for by the Bayesian model: as the posterior is proportional to the prior, the posterior mean is pulled towards the numbers that, under the prior, are more frequent. This shifts upwards the estimates in the larger-is-more-probable condition, in comparison to the smaller-is-more-probable condition (Fig. 3A). Hence the central tendency is not an arbitrary attraction towards the middle of the range (or towards the middle of the response slider), but it appears instead to result from the observer taking into account, in the choice of a response, the relative frequencies of different magnitudes.

In the stakes experiment, the prior is uniform and thus the frequencies of the different magnitudes are all equal. Thus the Bayesian posterior does not ‘favor’ small or large numbers, and one might predict that the central tendency should thus be the same in the two conditions of this experiment. But we find in this experiment a shift of the estimates very similar to that found in the priors experiment, with larger responses in the larger-has-higher-stakes condition than in the smaller-has-higher-stakes condition (Fig. 3B). Here also this modulation of the central tendency is reproduced by the Bayesian model (Fig. 4B). Bayesian responses are indeed pulled towards the numbers that result in higher rewards (while in the priors experiment, Bayesian responses are pulled towards the numbers that result in higher probabilities of reward). This behavior emphasizes the role of the objective (or loss) function in Bayesian decision theory, in addition to the prior. In fact, in the expression of the optimal Bayesian response (Eq. 2), the prior *p*(*x*) and the stakes function *q*(*x*) have interchangeable roles. Thus our manipulation of the stakes function modulates the behavior of the Bayesian observer in very much the same way that does our manipulation of the prior. The statistics of subjects’ responses are also impacted in remarkably similar ways by the two manipulations, and each in a fashion that is well captured by the Bayesian model (Figs. 3, 4). Our work adds to a literature that has previously exhibited the influence of reward on perceptual judgments [70–75], including in numerosity estimation [18]. (The paradigm of the stakes experiment however differs slightly from that employed in these studies, in which the reward for each trial is typically revealed to the subject before stimulus onset: in the stakes experiment, by contrast, the stimulus itself determines the size of the potential reward, and thus all trials have the same value *a priori* ). Our work shows how the influence of reward is similar to the influence of prior probabilities. As such, it substantiates the idea that perceptual decision-making and economic decision-making should be understood within a common framework [76].

Subjects however do not seem to perfectly learn the prior and the stakes function. Despite the symmetry of these functions in the two conditions of each experiment, the behavior of subjects in one condition does not perfectly mirror their behavior in the other condition, especially in the stakes experiment. We have noted that in the smaller-has-higher-stakes condition, although subjects shift the number where the MSE is minimized toward a lower number (as compared to the larger-has-higher-stakes condition), still the minimum is reached for a number above 60, i.e., not where numbers have high stakes (Fig. 3F). Our ‘subjective-asymmetric’ models capture this behavior, and suggest that subjects have not perfectly learned the stakes functions and the priors in these experiments. This impacted their performance: subjects collected significantly more points in the larger-has-higher-stakes condition than in the smaller-has-higher-stakes (p-value of t-test of equality: 0.038), resulting in a reward on average $0.24 higher in the former condition (average reward in the stakes experiment, including both conditions: $8.58). Although the effect is less pronounced in the priors experiment (where subjects seem to minimize the MSE toward smaller numbers, in the smaller-is-more-probable condition; Fig. 3E), it is in fact more consequential, as in this experiment the frequency of the resulting errors is increased. We find that subjects collected significantly more points in the larger-is-more-probable condition than in the smaller-is-more-probable (p-value of t-test of equality: 0.003), resulting in a reward on average $0.29 higher in the former condition (average reward in the priors experiment, including both conditions: $8.40). This emphasizes the importance of matching one’s beliefs with the true statistics, and the true stakes, of one’s environment.

The importance of the prior and the stakes raises the question as to where in the brain these quantities are encoded. There is currently no consensus regarding the neural representations of the prior. The theory of efficient coding mentioned above implies that the properties of sensory neural populations implicitly encode a long-term prior about the incoming stimuli [46]. But the prior seems to be also represented at higher levels of processing. In the macaque, the inferior temporal cortex, a later-stage visual processing area, appears to carry information about the statistics of stimuli [77]. An fMRI study of humans performing a random-dot motion task with varying priors finds that several frontoparietal regions are sensitive to the prior [78]. It might be the case that the prior is indeed encoded in many locations: a study using brain-wide data from the International Brain Laboratory concludes that in the mouse, the prior is encoded in 20% to 30% of brain regions, spanning all levels of processing: sensory, motor, and cortical regions [79]. The authors argue that this should allow for carrying out complex inferences in any direction, over a large Bayesian network.

Our theoretical framework suggests that the stakes have a role very similar to that of the prior (Eq. 2). Consistent with this idea and with the results just mentioned, the literature suggests that value-based information is also found at multiple levels in the brain. For instance, in the fMRI random-dot study mentioned above, the authors manipulate the payoff (in addition to the prior) and they find that similar frontoparietal areas are sensitive to these manipulations [78]. In early visual areas (like V1), value has been shown to increase response amplitude and to sharpen the tuning profiles of the relevant sensory populations, suggesting that sensory representations at a relatively early stage can be modulated by value [72, 73]. More traditionally, however, value representation in the brain is associated with the ‘reward system’, which comprises cortical and subcortical structures such as the ventromedial prefrontal cortex (vmPFC), the ventral striatum (VS), and the ventral tegmental area (VTA). In rhesus macaques, for instance, VS and vmPFC were shown to exhibit very similar neural sensitivities to value-related information, including the absolute value of an option, but also the probability of winning in a gambling task [80]. A possibility is that value (or expected value) is primarily represented in these areas, which then modulate neural sensitivities in sensory regions. Paralleling the question above on where the prior is encoded, the multiplicity of the locations of value signals in the brain is a source of debate in neuroeconomics and related fields [81]. As noted, some regions encoding information about value may also encode information about related probabilities. Adding to the complexity of the problem, different representations of probabilities and values across different locations in the brain may be inconsistent with each other [82].

Although our experimental results for the prior and the stakes experiment are both in line with the prediction of the Bayesian model, we have noted that the behavioral data suggest that the stakes functions might have been learned less accurately than the priors. A possibility is that these two functions are not learned the same way because they are not represented the same way in the brain. We also note however that the prior might be more salient: subjects are indeed more often exposed to the more probable numbers, in the priors experiment, but they are not more often exposed to the higher-stakes numbers, in the stakes experiment. If priors and values are not learned equivalently well, one may wonder which would yield a stronger effect if the two were conflicting, e.g., if some numbers had both high stakes and low probability, and conversely for other numbers. This would capture interesting (and not unusual) scenarios in which high-stakes events have low probabilities, and conversely. Our current results suggest that the prior may show a stronger influence, but we leave this question to further investigations. In any event, we have noted that in both experiments, model fitting suggests that subjects strongly underestimate the ratios characterizing the correct priors and stakes functions. We surmise that with more training, human subjects should be able to incorporate more accurate subjective functions in their decisions. This raises the question as to how priors and values are learned. In economics, a recent model of ‘adaptive utility’ proposes an iterative mechanism to construct a utility function optimally adapted to any distribution of reward, for an agent who has a limited representational capacity [83]. In psychology, studies on the ability of human subjects to learn priors are surprisingly scarce, and call for further experimental and theoretical investigations on prior learning [84, 85].

## Methods

### Details of the tasks

**Trials** Each condition of each task started with 15 ‘learning’ trials, in which the correct number of dots was shown alongside the cloud of dots. No response was required from the subject in these trials. The next 30 trials were ‘feedback’ trials, in which the subject was shown the correct number, after providing their estimate. These were followed by 120 ‘no-feedback’ trials, in which the correct number of dots was not shown, which we assumed would limit the risk of residual learning dynamics and of trial-to-trial sequential effects. All the analyses presented in this paper were conducted on the basis of the data obtained in the ‘no-feedback’ trials. The task was coded with jsPsych [86].

**Reward** In both experiments, the reward had two components: a fixed $3 USD base pay, and a performance bonus. The performance bonus was a function of the total number of points accumulated by the subject in the experiment. In the priors experiment, every 1000 points were worth 32¢. In each trial of the priors experiment, the number of points earned by the subject was a function of the difference between the correct number, *x*, and the subject’s provided response, *x*^, as 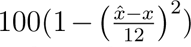. Subjects earned an average of $8.40 (s.d.: $2.10) in the priors experiment. In the stakes experiment, every 1000 points were worth 6¢. The reward in each trial of the stakes experiment depended on the stakes function, *q*(*x*), as 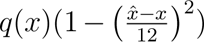. Subjects earned an average of $8.58 (s.d.: $1.61) in the stakes experiment.

**Subjects** Each subject participated in one experiment, and experienced the two conditions of this experiment (which were presented in counterbalanced orders). Subjects were recruited on Amazon Mechanical Turk through CloudResearch [87]. 120 subjects participated in the stakes experiment (66 male, 54 female; average age: 41.4, s.d.: 10.2), and 80 subjects participated in the priors experiment (47 male, 31 female, 2 non-binary; average age: 39.9, s.d.: 9.9). The study protocol was approved by the Institutional Review Board (IRB) of Harvard University (protocol IRB15-2048). As described in the preregistrations of the experiments, we excluded from the analysis the responses of all the subjects who obtained a performance bonus lower than $0.50. This resulted in the exclusion of 3.3% of subjects in the stakes experiment and 7.5% of subjects in the priors experiment.

### Data analysis

The statistics presented in Figure 3 correspond to the posterior-mean estimates of the fixed-effect components of a statistical model that included subject-specific random effects. In particular, the statistical model was specified by the three following equations:

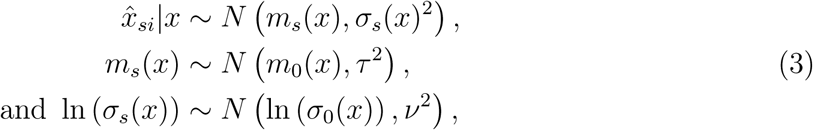

with the priors

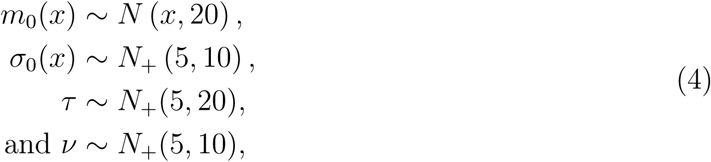

where *x*^_si_ was the response of subject *s* in trial *i*, and *N*_+_ is the Gaussian distribution truncated to the positive numbers. This statistical model was estimated using Stan with the HMC-NUTS sampler [88] (10 chains of 1000 samples each, following 1000 warmup iterations.) The shaded areas in Figure 3 correspond to the 5th and 95th percentile of the posterior. The stars indicate, for each quantity, that the Bayesian p-value is lower than 0.005, where the Bayesian p-value is defined as the posterior probability that the sign of the quantity’s difference across conditions is opposite to the sign apparent in the Figure. All other data analyzes were conducted using NumPy and Scipy, and figures were made using Matplotlib [89–91].

### Model fitting and model selection

We fit the nine models to the behavioral data by maximizing their likelihoods. In order to compute the likelihood function, i.e., the probability of a response *x*^ conditional on a presented number *x*, *p*(*x*^|*x*), we first compute the distribution of the optimal estimate, *x*^*^, conditional on *x*, *p*(*x*^*^|*x*), using the distribution of the internal signal, *r*, conditional on *x*, *p*(*r*|*x*), as specified by the model, and the expression of *x*^*^(*r*) given in Eq. 2. We note that the distribution of the Bayesian mean, with the priors and the stakes functions that we use in our experiment, does not belong to a canonical family of distributions, and neither its density nor its cumulative distribution function admit an analytic expression. We thus resort to numerical computations. Second, we compute (numerically) the distribution of estimates *x*^ conditional on the optimal estimate, *x*^*^, *p*(*x*^|*x*^*^), which is a normal distribution centered on *x*^*^ and clipped to the interval 41-80. Finally we compute *p*(*x*^|*x*) on the basis of the two distributions just presented, as 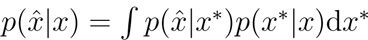.

For each model we fit each subject’s dataset separately (with a different set of parameters per subject). The resulting BICs are reported in Table 1. To obtain a more refined view on the relative prevalence of each model in the population of subjects, we conducted a ‘Bayesian model selection’ analysis [34, 35]. This procedure enabled the derivation of a Bayesian posterior over the nine models, as a Dirichlet distribution. From this posterior we computed, first, the expected probability of each model: this is the expected value of the probability that the behavior of a subject chosen randomly in the population follows the given model. On the basis of the posterior we also computed the protected exceedance probability (PXP) of each model, defined as the probability that the model is the most prevalent in the population, taking into account the possibility that differences in model evidence may be due to chance [35]. The estimation of the PXP was derived from the ‘exceedance probability’, which we estimated by sampling 10 million times the Dirichlet posterior, and counting the number of times each model has the largest probability. The expected probabilities and the PXPs are reported in Table 1. We report here the sum of the parameters of the Dirichlet posterior, which indicates the concentration of the posterior: for the priors experiment the sum is 83; for the stakes experiment the sum is 125.

### Preregistrations

The two experiments were preregistered on AsPredicted. The preregistration for the priors experiment can be found here: https://aspredicted.org/hb3s4.pdf. The preregistration for the stakes experiment can be found here: https://aspredicted.org/6sg5b.pdf.

## Code and data availability

The experimental code, the data, and the data-analysis code pertaining to the presented results are available here: https://osf.io/9ueza/?view_only=d85803890cdf42088c3df563096fdbba.

## Acknowledgments

We thank Peng Qian for helpful conversations. This research was supported by the National Science Foundation (DRL-2024462) and the Air Force Office of Scientific Research (FA9550-20-1-0413).

